# Selected humanization of yeast U1 snRNP leads to global suppression of pre-mRNA splicing and mitochondrial dysfunction in the budding yeast

**DOI:** 10.1101/2023.12.15.571893

**Authors:** Subbaiah Chalivendra, Shasha Shi, Xueni Li, Zhiling Kuang, Joseph Giovinazzo, Lingdi Zhang, John Rossi, Anthony J. Saviola, Jingxin Wang, Robb Welty, Shiheng Liu, Katherine Vaeth, Z. Hong Zhou, Kirk C. Hansen, J. Matthew Taliaferro, Rui Zhao

## Abstract

The recognition of 5’ splice site (5’ ss) is one of the earliest steps of pre-mRNA splicing. To better understand the mechanism and regulation of 5’ ss recognition, we selectively humanized components of the yeast U1 snRNP to reveal the function of these components in 5’ ss recognition and splicing. We targeted U1C and Luc7, two proteins that interact with and stabilize the yeast U1 (yU1) snRNA and the 5’ ss RNA duplex. We replaced the Zinc-Finger (ZnF) domain of yU1C with its human counterpart, which resulted in cold-sensitive growth phenotype and moderate splicing defects. Next, we added an auxin-inducible degron to yLuc7 protein and found that Luc7-depleted yU1 snRNP resulted in the concomitant loss of PRP40 and Snu71 (two other essential yeast U1 snRNP proteins), and further biochemical analyses suggest a model of how these three proteins interact with each other in the U1 snRNP. The loss of these proteins resulted in a significant growth retardation accompanied by a global suppression of pre-mRNA splicing. The splicing suppression led to mitochondrial dysfunction as revealed by a release of Fe^2+^ into the growth medium and an induction of mitochondrial reactive oxygen species. Together, these observations indicate that the human U1C ZnF can substitute that of yeast, Luc7 is essential for the incorporation of the Luc7-Prp40-Snu71 trimer into yeast U1 snRNP, and splicing plays a major role in the regulation of mitochondria function in yeast.

## Introduction

Gene regulatory mechanisms have played a predominant role in the evolution of organismal diversity, including the origin of multicellularity (King and Wilson 1975; Carroll 2005; Wittkopp and Kalay 2011; Kianianmomeni 2015). Transcriptional and posttranscriptional processes constitute the early steps of gene expression. In eukaryotes, pre-mRNA splicing is a major gene regulatory mechanism (Girardini et al. 2023). Only 2.8% of the human genome represents coding sequences, while 35% corresponds to introns (Hatje et al. 2019). The intronic expansion has resulted in a huge transcriptomic/proteomic diversity, as 90% of human genes are alternatively spliced (Calarco et al. 2007). While tissue- and development-specific regulation of splicing facilitated organismal complexity, it necessitated the evolution of the spliceosome itself, to keep pace with the intronic and species diversity even among microbes (Sales-Lee et al. 2021). The spliceosome is comprised of five highly conserved uridylyl-rich small RNA-protein complexes (U snRNPs), namely, U1, U2, U4, U5 and U6 (Wahl et al. 2009). U1 snRNP (U1) is critical for the recognition of introns, specifically, the 5’ splice site (5’ ss) and forming the first commitment complex of the splicing cycle (E-complex in humans and CC1/CC2 in yeast). The preponderant role of U1 in gene expression is beyond pre-mRNA splicing, including upstream events of gene expression. In mammals, U1 is critical for enhancing transcription initiation (Damgaard et al. 2008; Engreitz et al. 2014; Rose 2018), rate of transcription elongation (Mimoso and Adelman 2023), determining promoter directionality (Almada et al. 2013) and, suppressing premature polyadenylation and transcriptional termination (Kaida et al. 2010; Di et al. 2019). Many human genetic diseases involving splicing defects related to the failure of 5’ ss recognition by the human U1 snRNP (hU1) (Lorson et al. 1999; Roca et al. 2008; Juschke et al. 2021). A detailed structural and functional analysis of the hU1 is critical for understanding splicing regulation and implications for human health.

Human U1 snRNP often has transient and contextual interaction with alternative splicing factors, many of whose identities and roles are not fully known. These intricate genetic and molecular interactions may not be easily studied *in vitro*, cell culture or other metazoan model systems. Yeast is a facile model to address outstanding questions in splicing regulation, owing to its simplicity and versatility, in addition to the conservation of splicing machinery (Fabrizio et al. 2009; Meyer and Vilardell 2009). As expression of human homologs can complement yeast genes of diverse pathways, yeast has been successfully used to dissect complex biological pathways relevant to human physiology and pathology (Garge et al. 2020; Boonekamp et al. 2022; Hunter et al. 2023) as well as drug discovery (dos Santos and Sa-Correia 2009; Zimmermann et al. 2018). Yeast has fewer (334; ∼5% of the genome) and smaller introns with a highly conserved 5’ ss, and most of them are co-transcriptionally processed (Spingola et al. 1999; Aslanzadeh et al. 2018; Talkish et al. 2019). Being a unicellular eukaryote uncomplicated by tissue type diversity, gene regulation in yeast is tightly linked to the phenotype (Airoldi et al. 2009; Strassburg et al. 2010; Tamari et al. 2014). In turn, gene expression itself is closely aligned to pre-mRNA splicing due to an enrichment of introns in ribosomal protein genes (RPGs) and the high transcription rate of intron-containing genes (both RPGs and non-RPGs) in yeast (Ares et al. 1999; Juneau et al. 2006; Lukacisin et al. 2022). This permits reliable correlation of any introduced alterations in the splicing machinery with changes in molecular, supramolecular, and cellular/organismal phenotypes.

Human U1 snRNP contains a 164 nt U1 snRNA that threads through the heptameric ring formed by seven Sm proteins, and three additional proteins U1A, U1C, and U1-70K. Although earlier studies suggested that U1C makes important contribution to the sequence-specific recognition of 5’ ss (Du and Rosbash 2002), structural analysis of hU1 snRNP revealed that 5’ ss is primarily recognized via RNA-RNA binding to the 5’ end of U1 snRNA (Kondo et al. 2015). U1C uses its ZnF domain to bind the sugar phosphate backbone but not the bases of the 5’ss-U1 snRNA duplex and is required for the duplex stability, particularly when 5’ ss sequences are non-canonical (Kondo et al. 2015). The 5’ end of U1 snRNA that base-pairs with 5’ ss sequence is invariant across all eukaryotes, from yeast to human (Kretzner et al. 1987). However, human 5’ ss sequences are highly divergent, unlike the predominantly uniform 5’ ss sequence of yeast introns. It follows that the duplex-stabilizing role of U1C becomes especially relevant in the case of hU1 snRNP.

Yeast U1 snRNP contains a much larger snRNA, homologs of all human U1 snRNP proteins, and seven additional stably associated proteins. Cryogenic electron microscopy (cryoEM structure of the yeast U1 snRNP demonstrated that the yeast U1 snRNP core (composed of U1 snRNA and homologs of the human U1 snRNP proteins) is highly similar to the entire human U1 snRNP, with additional yeast auxiliary proteins stably bound to the core (Li et al. 2017). Of these auxiliary yeast proteins, Luc7 also uses a ZnF domain to bind the 5’ss-U1 snRNA duplex, presumably providing additional stability for the duplex. There are three human homologs (Luc7L, Luc7L2, Luc7L3) of Luc7 that are known alternative splicing factors (Puig et al. 2007). The human Luc7Ls bind U1 snRNA as well as an AAGAAG sequence in the exon near weak 5’ ss (Daniels et al. 2021), suggesting that they may also help stabilize the 5’ ss-U1 snRNA duplex similar to yeast Luc7. In the current study, we have made selected modifications (humanization) of yeast U1 snRNP components to unravel the mechanistic details of 5’ ss recognition of splicing regulation in eukaryotes. Our studies reveal novel details on structure-function relationships of two essential U1 snRNP proteins and how these molecular interactions are integrated into organismal responses to external and internal stimuli.

## Results

### Humanization of yU1C leads to moderate splicing defects

The N-terminal region of U1C containing the ZnF domain is highly conserved between human and yeast, both in sequence (50% identity and 69% similarity) and its role in forming the first commitment or the E complex (Nelissen et al. 1991; Muto et al. 2004; Schwer and Shuman 2014; Kondo et al. 2015; Schwer and Shuman 2015), while its C-terminal region is highly divergent (**Fig. 1A**). We introduced the full length human U1C into a yeast U1C shuffle strain (Schwer and Shuman 2014) and showed that human U1C cannot substitute yeast U1C (**Fig. 1B**). We next replaced the first 36 amino acids of yeast U1C with that of human (abbreviated as h-yU1C for humanized yeast U1C) using CRISPR in a yeast strain where U1A is TAP tagged with protein A and Calmodulin Binding Peptide (CBP) (Li et al. 2017). h-yU1C grew similarly as the wild-type (WT) yeast strain at 30 °C and 37 °C, although it grew slower at low temperature (23 °C) and even more so at 17 °C (**Fig. 1C**).

**Figure 1.**
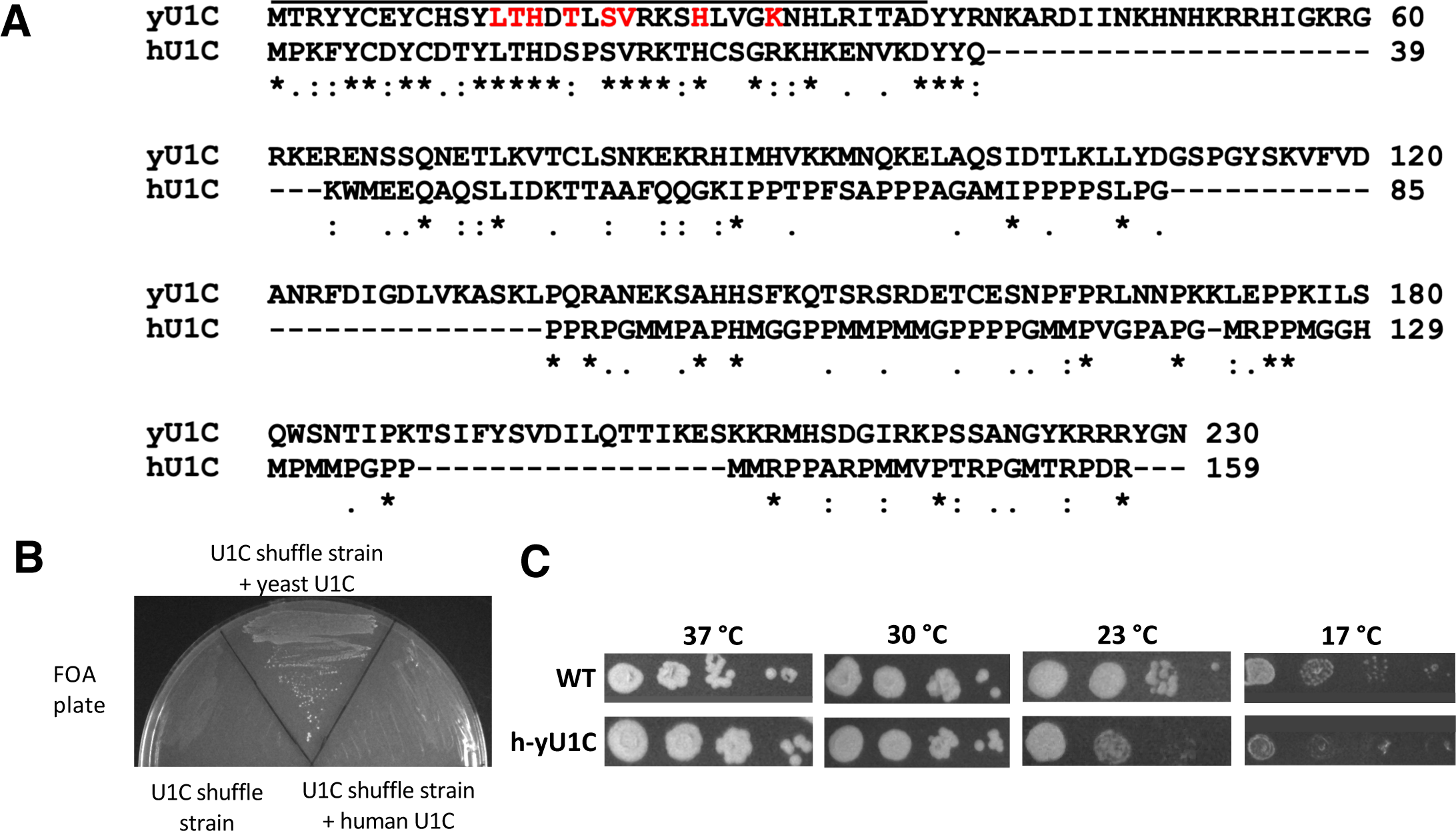
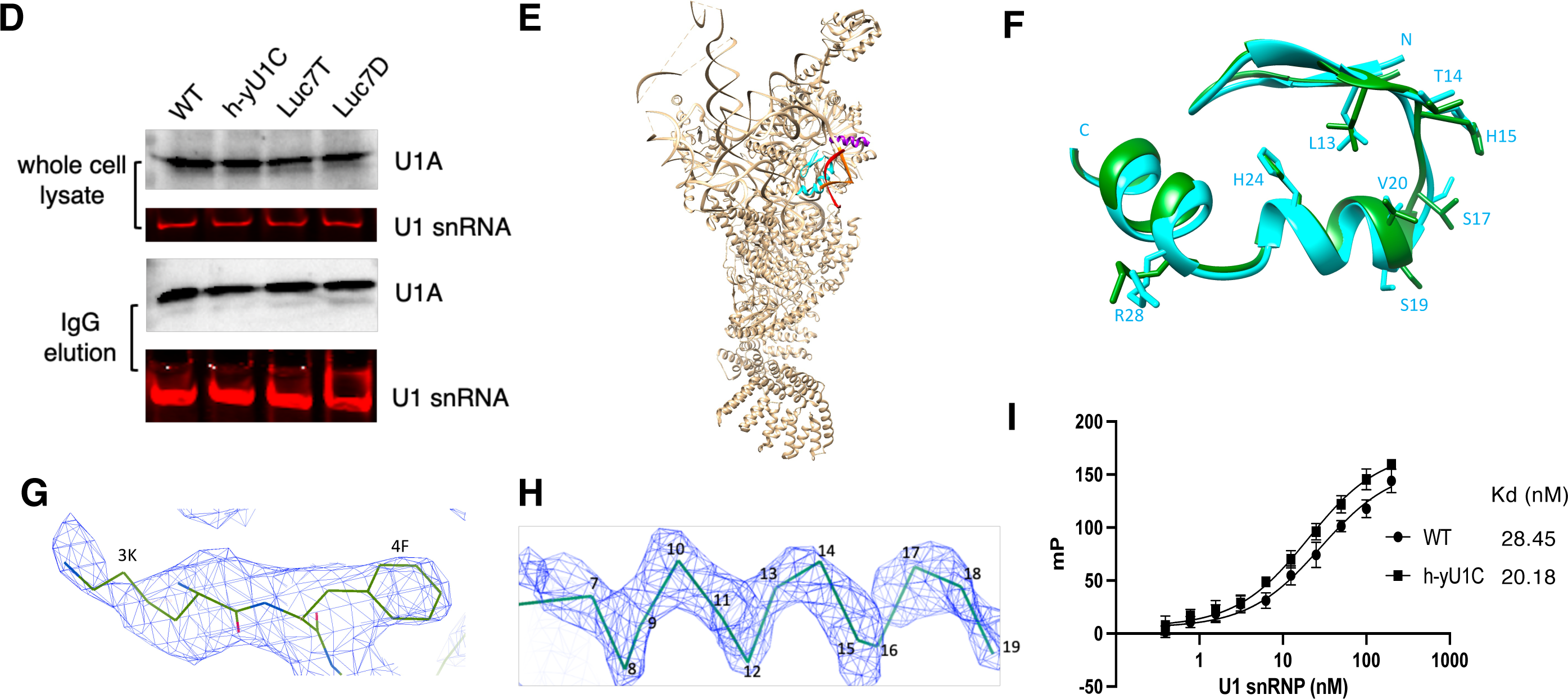
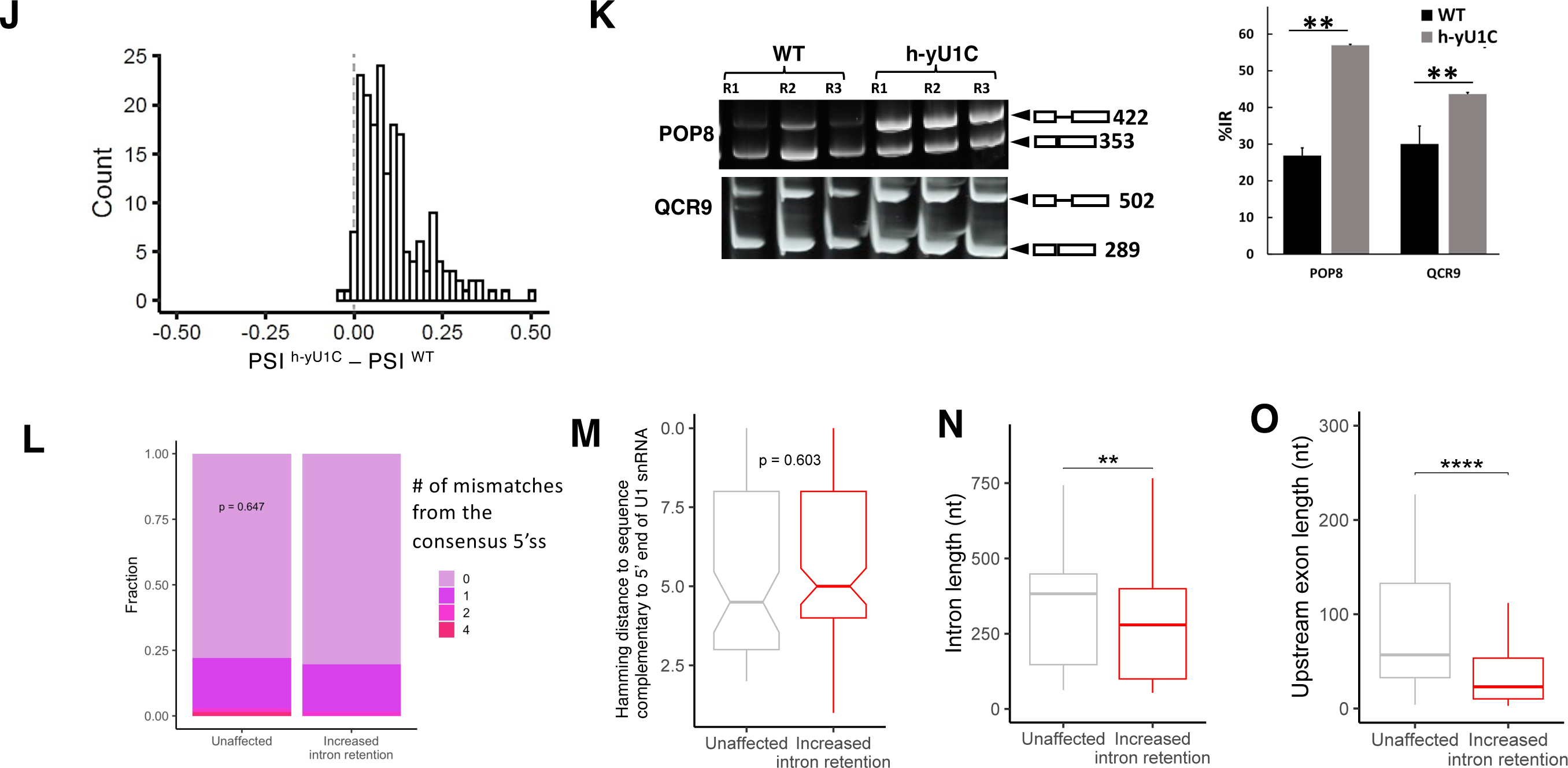
Humanization of yU1C leads to moderate splicing defects. **(A)** Yeast and human U1C are highly conserved in the ZnF domain as demonstrated by sequence alignment using Clustal Omega (Sievers et al. 2011). * = identical residues, : = conserved residues,. = Less conserved residues, -- = Gaps in one of the pairs. The line above the sequence indicates the yeast ZnF domain that was replaced with the corresponding human sequences. Red font indicates residues interacting with the U1 – 5’ ss RNA duplex in the cryoEM structure of the yeast E complex (Li et al. 2019). **(B)** A yeast U1C shuffle strain (Schwer and Shuman 2014) alone, transformed with yeast U1C, or transformed with human U1C were plated on FOA plate. Only the U1C shuffle strain transformed with yeast U1C grew. **(C)** Yeast strain with humanized yeast U1C is cold-sensitive as demonstrated by spot assay with 10x serial dilutions at different temperatures. **(D)** U1A (probed by an anti-CBP antibody) and U1 snRNA (probed by solution hybridization) levels in whole cell lysate and after IgG pull down using the TAP-tag on U1A are similar between the WT and h-yU1C strain at both 30 and 17 °C. **(E)** Overall structure of the U1 snRNP ^h-yU1C^. The ZnF domain of U1C that was humanized is shown in cyan, the 5’ ss in red, the first 10 nt of U1 snRNA in orange, and the N-terminal helix of Luc7 in purple. **(F)** Superimposition of the ZnF domain of WT (green) and h-yU1C (cyan) backbone. Residues in h- yU1C that interact with the U1 – 5’ ss RNA duplex are shown in stick model and labeled. **(G)** An example of residues that are different between human (Lys3 and Phe4) and yeast (Arg3 and Tyr4) with their corresponding cryoEM density map (mesh) and atomic model (sticks) in the U1 snRNP ^h-yU1C^ structure. **(H)** CryoEM density map (mesh) surrounding the N-terminal helix of Luc7 (backbone model in sticks) in the U1 snRNP ^h-yU1C^ structure. **(I)** U1 snRNP ^h-yU1C^ has slightly lower Kd to an oligo containing the yeast canonical 5’ ss (GAUUCUG/GUAUGUUC where / designates the exon/intron boundary) than the WT U1 snRNP as determined by fluorescence polarization experiments. Fluorescence polarization values are baseline (oligo alone) subtracted to better align and compare the two binding curves. **(J)** RNAseq experiments revealed that humanized yeast U1C leads to poorer splicing of a moderate number of intron-containing genes as demonstrated by their increased PSIs (percentage spliced in, or percentage intron retention). **(K)** RT-PCR of POP8 and QCR9 show that splicing is impaired in the h-yU1C strain at 17 °C. POP8 and QCR9 were picked as examples of genes affected by humanizing yeast U1C in the RNAseq data as two of the most affected genes. **(L)** There is no significant difference between unaffected and retained introns in how close their 5’ splice sites are to the consensus sequence. p values were calculated using a binomial test considering the fraction of introns with perfect and imperfect matches to the consensus sequence of GTATGT. **(M)** There is no significant difference between unaffected and retained introns in how close their 5’ splice sites are to the sequence that complements the 10 nt at the 5’ end of U1 snRNA. p values were calculated using a Wilcoxon rank sum test considering the Hamming distances of introns to the sequence (AGGTAAGTAT) complementary to the 5’ end of U1 snRNA. **(N)** Increased intron retention significantly correlates with shorter intron length. **(O)** Increased intron retention significantly correlates with shorter upstream exon. p values were calculated using a Wilcoxon rank sum test in panels M and N. **, p<0.01. ****, p<0.0001.

We pulled down U1 snRNP using the protein A tag on U1A and demonstrated that the h-yU1C strain has similar levels of U1 snRNP as the WT strain at both 30 and 17 °C, by detecting the level of U1 snRNA using solution hybridization (Li and Brow 1993) and the level of U1A through its CBP tag by immuno-blotting (**Fig. 1D**). We purified U1 snRNP from the h-yU1C strain using the TAP tag on U1A and determined its cryoEM structure in complex with the Act1 pre-mRNA and BBP/Mud2 to 3.5 Å resolution (**Supplementary** Fig. 1), similar to what we have done with the WT yeast U1 snRNP (Li et al. 2019) (**Fig. 1E**). The structure demonstrates that h-yU1C binds to the U1 snRNA and 5’ ss duplex similarly to the WT U1 snRNP (**Fig. 1E**). The h-yU1C and yU1C main chains can be well superimposed (**Fig. 1F**), and the side chains of residues different between the two have clearly different densities as expected (**Fig. 1G**). Residues that contact the U1 snRNA and 5’ ss RNA duplex are mainly concentrated on the α-helix (residues 13-28) in the ZnF and have minimal changes in both sequence (**Fig. 1A**, T17 to S and K28 to R from yeast to human) and structure (**Fig. 1F**) between the two species. In addition, Luc7 is present in the structure with well-defined density for its N-terminal alpha helix that interacts with the Sm ring (**Fig. 1H**), although the rest of the Luc7 density is not as well defined as in the WT spliceosomal E complex (Li et al. 2019), potentially due to the much smaller particle numbers in the U1 snRNP ^h-yU1C^ data set.

We then evaluated the binding affinity between U1 snRNP ^h-yU1C^ and the 5’ ss pre-mRNA in comparison to the WT U1 snRNP, using fluorescence polarization experiment and a 15nt RNA (GAUUCUG/GUAUGUUC where / denotes the exon/intron boundary) containing the consensus 5’ ss. We found that the Kd between U1 snRNP ^h-yU1C^ and the 5’ ss RNA is 20.18 nM (18.11 – 22.50 nM at 95% confidence interval), slightly lower than the Kd between WT U1 snRNP and the same 5’ ss oligo which has a Kd of 28.45 nM (23.97 – 33.85 nM at 95% confidence interval) (**Fig. 1I**).

We carried out RNA-seq analyses of the h-yU1C and WT strain, from three biological replicates of each. These analyses consistently showed that a number of intron-containing genes in the h-yU1C strain have greater intron retention (IR), measured as the percentage of spliced in (PSI) values, than the WT (**Fig. 1J**). Of the 190 yeast intron-containing genes with sufficient reads, ∼17% (32 out of 190) showed 20% or higher intron retention, and ∼43% (83 out of 190) had 10% or higher intron retention in the h-yU1C strain compared to the WT (**Fig. 1J**). As expected, similar intron retention was observed at lower temperature (**Fig. 1K**). The intron retention does not correlate with whether the intron contains the canonical yeast 5’ ss (AG/GUAUGU where / designates the exon/intron boundary), or whether the intron perfectly complements the 5’ end of U1 snRNA (note that the yeast canonical 5’ ss sequence AG/GUAUGU does not form a perfect complement to the 5’ end of U1 snRNA which has the sequence of ACUUACCU) (Spingola et al. 1999) (**Fig. 1L, M**). On the other hand, intron retention was greater for genes with shorter intron length or shorter upstream exon (**Fig. 1N, O**).

### Luc7 triple mutant has minimal effect on splicing

Unlike yeast Luc7 which is an integral component of yU1 snRNP, its three human homologs Luc7L, Luc7L2, and Luc7L3 (**Fig. 2A**) bind only transiently to hU1 snRNP in a tissue- and development-specific manner (Howell et al. 2007). We first attempted to humanize yU1 snRNP by replacing yLuc7 with its human homologs in a Luc7 shuffle strain. However, none of the shuffle strain transformed with Luc7Ls grew on 5-fluoroorotic acid (FOA; **Fig. 2B**), indicating that hLuc7Ls are unable to replace Luc7 in yeast.

**Figure 2.**
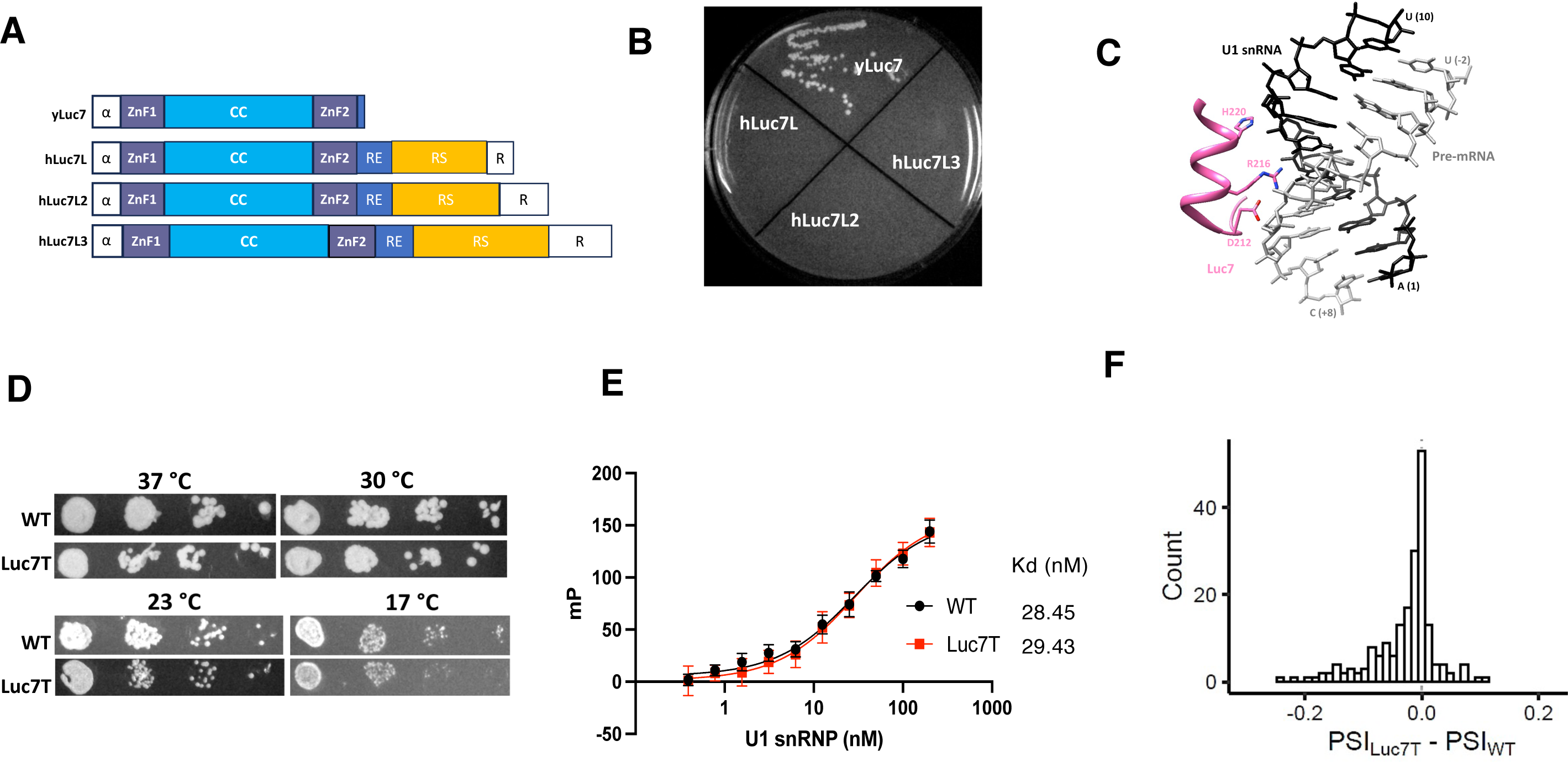
Luc7 triple mutant has minimal effect on splicing. **(A**) Domain organization of the yeast and human Luc7 homologs. α represents alpha helix. CC represents a colied-coil region. RE, RS, and R represent Arg/Glu, Arg/Ser, and Arg-rich regions. **(B)** Human Luc7 homologs cannot replace the yeast Luc7, as demonstrated by the lack of growth on a FOA plate after hLuc7Ls were introduced into the yeast Luc7 shuffle strain. **(C)** CryoEM structure of the yeast E complex (Li et al. 2019) indicates that residues D212, R216, and H220 are in close proximity of the U1 – 5’ ss RNA duplex. **(D)** Spot assay with 10x serial dilution shows Luc7T has similar growth as the parental WT strain, with a very mild growth retardation at 17 °C. **(E)** Fluorescence polarization experiments demonstrated that U1 snRNP ^Luc7T^ and U1 snRNP ^WT^ have indistinguishable binding affinity to the 5’ ss oligo. Fluorescence polarization values are baseline (oligo alone) subtracted to better align and compare the two binding curves. **(F)** Luc7T has minimal effect on PSIs of intron-containing genes in both the positive and negative direction as revealed by RNAseq experiments.

The yeast E complex structure we determined (Li et al. 2019) had clear density for the backbone of ZnF2 of yLuc7. However, the side chain density was often missing, making it difficult to unambiguously model the side chain conformation. Residues D212, R216, and H220 are in close proximity of the U1 snRNA and 5’ ss duplex (**Fig. 2C**) and these residues potentially interact with the RNA duplex. Previous genetic experiments also showed that the D216A mutant affects splicing of the Sus1 gene which has a weak non-canonical 5’ ss (Agarwal et al. 2016). Since we could not replace yLuc7 with the entire hLuc7Ls, we generated a Luc7 triple mutant (abbreviated as Luc7T) which contains the D212A/R216A/H220C substitutions (we mutated H220 to a Cys instead of Ala to maintain the Zinc Finger) with the objective of weakening the yLuc7 interaction with the 5’ ss.

Yeast strain carrying Luc7T showed no substantial growth defects except for a very subtle growth retardation at 17 °C (**Fig. 2D**). We purified U1 snRNP ^Luc7T^ from yeast using the TAP tag on U1A and determined its binding affinity to the 5’ ss oligo (Kd = 29.43 nM, 25.95 – 33.42 nM at 95% confidence interval). This Kd was indistinguishable from that of the U1 snRNP ^WT^ and the same oligo (**Fig. 2E**), suggesting that the three Luc7 residues (D212, R216, and H220) are not critical for binding to the U1 – 5’ ss RNA duplex.

Consistent with this observation, our RNA-seq analysis of the Luc7T strain revealed only minor splicing difference compared to the WT. Of the 193 yeast intron-containing genes with sufficient reads, only 9 displayed a PSI decrease of 10-19% and 2 with a PSI decrease of 20-25% (less IR or improved splicing) in the Luc7T strain compared to the WT (**Fig. 2F**).

### Luc7 depletion leads to the loss of Prp40 and Snu71 in U1 snRNP, and extensive splicing defects

Given that the Luc7T mutant did not significantly interrupt its interaction with the 5 ss – U1 snRNA duplex, we introduced an auxin-inducible degradation (AID) system (Mendoza-Ochoa et al. 2019) in the h-yU1C strain to deplete Luc7 (an essential protein in yeast) upon auxin addition. We fused an AID-tag to the C-terminus of Luc7 and co-expressed TIR1, an auxin-binding receptor, under the control of a β-estradiol inducible promoter. The addition of IAA (a common auxin) and β-estradiol leads to the expression and activation of TIR1, which in trun targets AID-tagged Luc7 for proteosome degradation. For simplicity, we designated the resulting strain as Luc7D (for Luc7 degron-tagged), although this strain also has the ZnF domain of yeast U1C humanized as the h-yU1C strain. After 6 hours of β-estradiol and IAA induction, we were able to completely deplete AID-tagged Luc7 (**Fig. 3A**). The h-yU1 snRNP strain showed a slow growth phenotype in the rich YPD medium (**Fig. 3B**).

**Figure 3.**
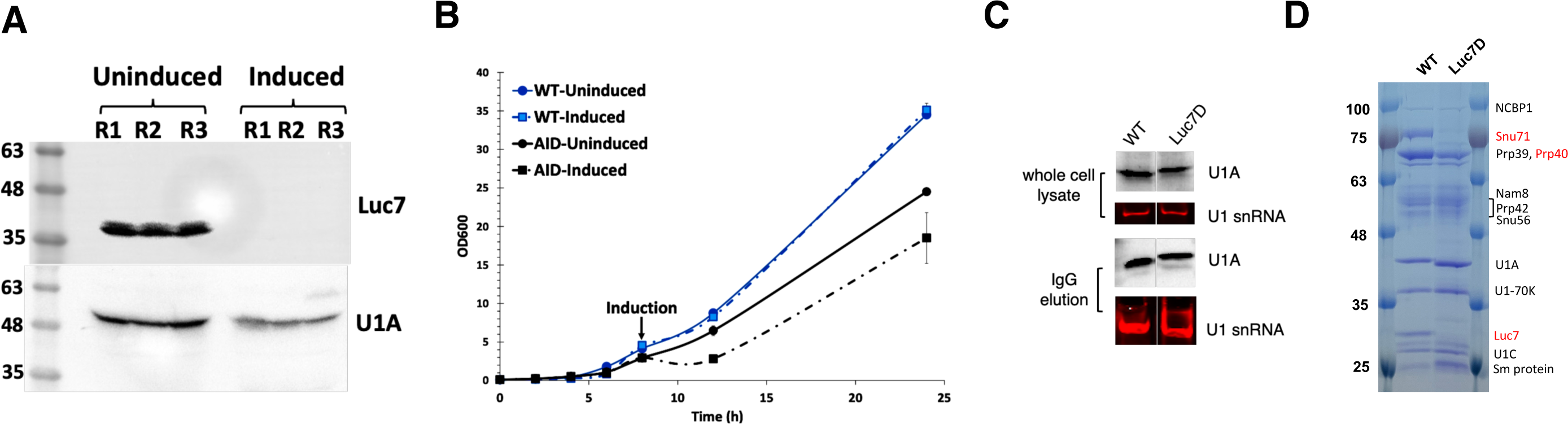
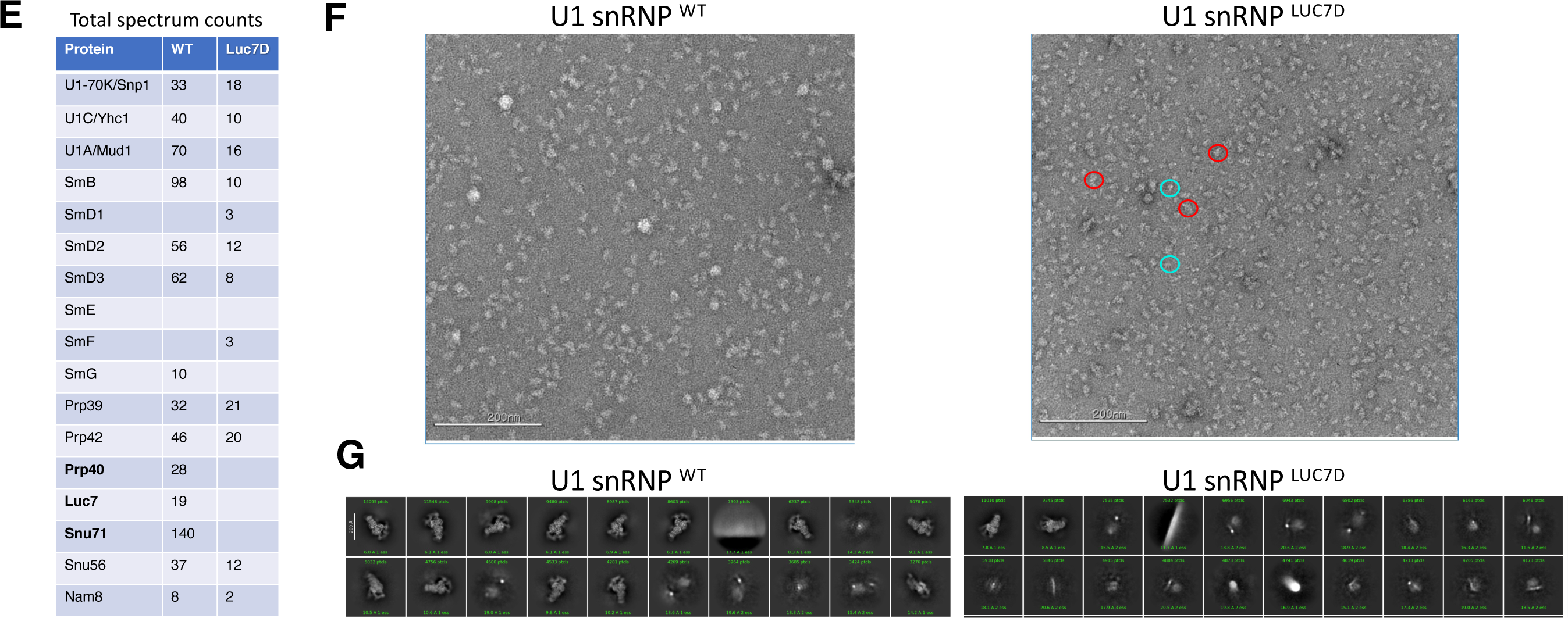
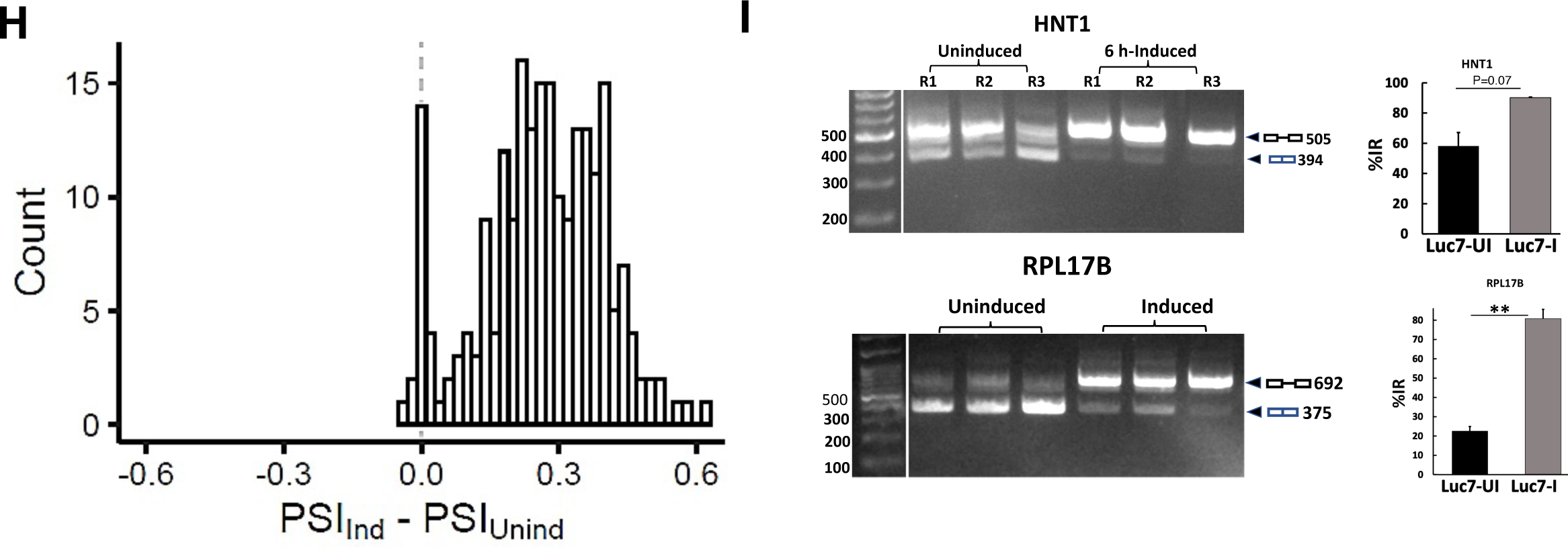
Luc7 depletion leads to the loss of Prp40 and Snu71 in U1 snRNP, and extensive splicing defects. **(A**) Luc7 was depleted after 6 hours of β-estradiol and IAA induction, as shown in Western blot of whole cell lysate using antibodies against the HA tag on Luc7 and CBP-tag on U1A. R1-3 represent three biological replicates. **(B)** The growth of Luc7D strain was retarded upon induction of Luc7 depletion, relative to the uninduced culture or the WT. **(C)** U1A (probed by an anti-CBP antibody) and U1 snRNA (probed by solution hybridization) levels in whole cell lysate and after IgG pull down using the TAP-tag on U1A are similar between the WT and Luc7D strains. **(D)** Coomassie stained SDS PAGE gel shows that purified U1 snRNP ^Luc7D^ after 6 hr induction is depleted of not only Luc7, but also Snu71 and Prp40 (indicated by the red font), compared to the WT U1 snRNP. **(E)** Mass spectrometry analyses of purified U1 snRNP ^Luc7D^ confirm the depletion of Luc7, Snu71, and Prp40 (bold font), compared to the WT U1 snRNP. Sm E, F, G proteins are small and more difficult to detect in mass spectrometry which led to the lack of peptides in both the WT and Luc7D samples. **(F)** Negative stain images demonstrate more smaller (likely broken down) particles in purified U1 snRNP ^Luc7D^ as compared to the WT. Red and blue circles highlight some intact and small particles. **(G)** The top 20 class averages (sorted by number of particles in each class) from 2D classification (Punjani et al. 2017) of a small cryoEM dataset show fewer classes and lower number of intact U1 snRNP particles in purified U1 snRNP ^Luc7D^ as compared to the WT. **(H)** Luc7 depletion after induction leads to massive splicing defects (manifested as increased PSI) in 203 out of 204 intron-containing genes with sufficient reads in RNAseq. **(I)** Examples of two genes whose splicing are significantly impaired upon Luc7 depletion as shown by RT-PCR products analyzed by native PAGE.

Luc7 depletion did not significantly affect the level of U1 snRNP, as revealed by the U1A and U1 snRNA levels in the IgG pull-down fraction (using the TAP-tag on U1A) of the cell extract from the Luc7D strain (**Fig. 3C**). We successfully purified U1 snRNP ^Luc7D^. Gel electrophoresis analysis showed that the complex was depleted in Luc7 as well as Prp40 and Snu71 (**Fig. 3D**), which is further confirmed by mass spectrometry (**Fig. 3E**). Negative stain image demonstrates that Luc7D can form U1 snRNP complex, although many particles are significantly smaller than the WT U1 snRNP, likely due to the instability of an incomplete U1 snRNP (**Fig. 3F, top**). 2D classification of a small cryo dataset also showed fewer classes of intact particles in the U1 snRNP ^Luc7D^ sample (**Fig. 3G, bottom**), indicating that U1 snRNP ^Luc7D^ is less stable than the WT U1 snRNP.

RNA-seq analysis showed that pre-mRNA splicing was severely impacted in the Luc7D strain after Luc7 depletion compared to the uninduced control (**Fig. 3H, I)**. Of the 204 intron-containing yeast genes with sufficient read coverage and consistent splicing changes in all triplicates, 203 demonstrated intron retention, over 96% of which showed > 10% IR increase compared to the uninduced sample.

### Luc7, Prp40, and Snu71 interact with each other through distinct domains

The loss of Prp40 and Snu71 upon Luc7 depletion prompted us to further investigate the interactions among Prp40, Snu71, and Luc7. Previous biochemical studies demonstrated that Prp40 and Snu71 can form a stable dimer, and Prp40, Snu71 and Luc7 form a trimer (Ester and Uetz 2008; Gornemann et al. 2011; Li et al. 2017). EM structures and biochemical studies also indicated that Luc7 uses its coiled-coil (CC) domain to bind Snu71 (residues 260-312 in the pre-A complex structure) (Li et al. 2019; Zhang et al. 2021). A boomerang shaped density near U1-70K in the yeast E complex structure was tentatively modeled as FF4-5 domains of Prp40 (Li et al. 2019). A longer stretch of density was observed in similar areas in the yeast pre-A complex structrure and was modeled as Prp40 FF1-6 (FF6 contacts U1-70K and FF1 contacts Luc7) based on the shape of the density and crosslinking and mass spectrometry data (Zhang et al. 2021). However, the model is not definitive due to the limited resolution of the density. For example, FF1 and FF2 are modeled right next to each other (close to the CC domain of Luc7) with no room for the long helix conncting the two domains (Zhang et al. 2021). Crosslinking and mass spectrometry of the pre-A complex also indicated that the C-terminal region of Snu71 interacts with FF2 and 3 of Prp40 (Zhang et al. 2021).

We further explored the interaction between Prp40 and Snu71 with an orthogonal approach using pull-down assays and purified GST-fused Prp40 domains and Snu71. We demonstrated that the Prp40 FF4 domain interacts directly and strongly with Snu71 (**Fig. 4A**). Purified FF2 and 3 domains are severely degraded and the lack of interaction observed in pull down is inconclusive. Using similar pull-down assays, we showed that the Snu71 region containing residues 382-537 directly interacts with the Prp40 FF4 domain (**Fig. 4B**). We further demonstrated that the WW domains (residues 1-75) of Prp40 directly interacts with Branch Binding Protein (BBP) in pull-down assays (**Fig. 4C**, pull-down results summarized in **Fig. 4D**). This observation is consistent with previous chemical shift perturbation experiments indicating that the PSPPPVYDA peptide (residues 94-102) in the N-terminal domain of BBP binds directly to the WW domains of Prp40 (Abovich and Rosbash 1997; Wiesner et al. 2002).

**Figure 4.**
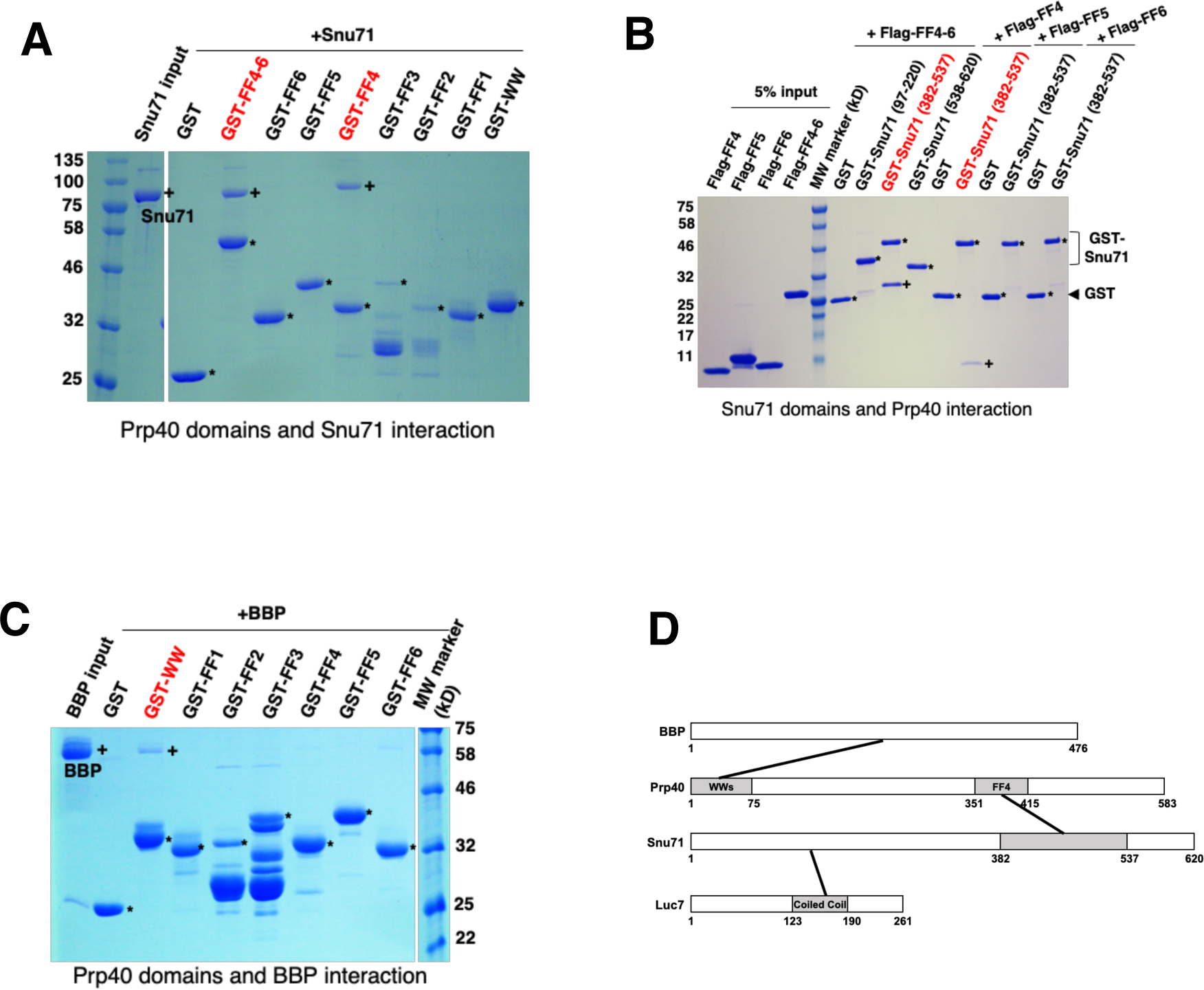
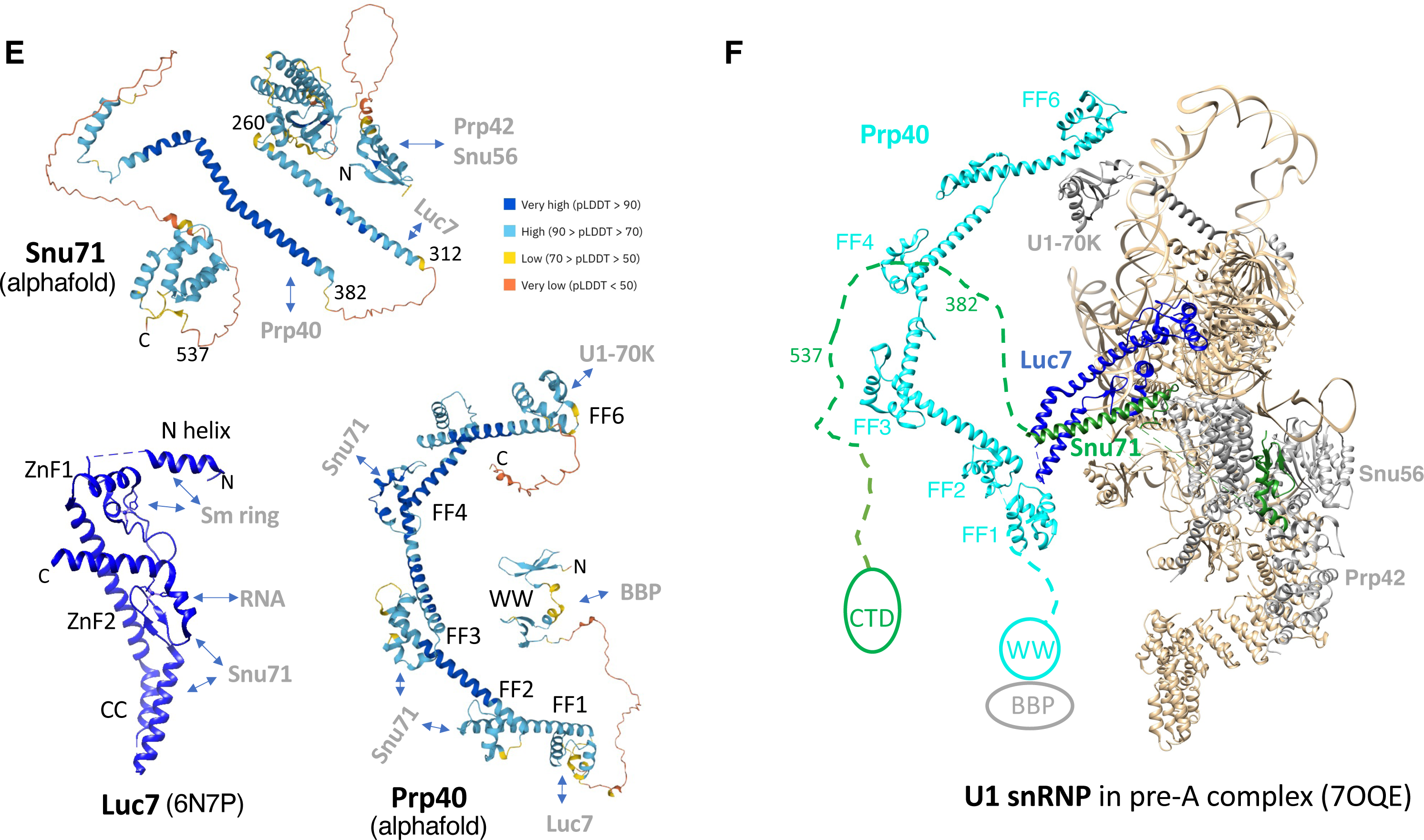
Probing the Prp40, Snu71, and BBP interaction using GST pull-down assays. **(A)** GST pull-down assays probing the interaction between Prp40 domains and Snu71. Various GST-fused Prp40 domains expressed and purified from *E. coli* (WW including both WW1 and 2, FF1, FF2, FF3, FF4, FF5, FF6, FF4-6) are used to pull down Snu71. Pull-down samples after extensive wash are visualized by SDS PAGE and Coomassie staining. “*” denotes bands corresponding to various GST-fusion Prp40 domains (GST-FF2 and FF3 have significant degradation). “+” denotes bands corresponding to Snu71. Red labels indicate Prp40 domains that pulled down Snu71. **(B)** GST pull-down assays probing the interaction between Snu71 domains and Prp40. Various GST-fused Snu71 domains (residues 97-220, 382-537, 538-620 while domains covering other regions of Snu71 cannot be expressed in *E. coli*) are used to pull down Flag-tagged Prp40 FF4, FF5, FF6, and FF4-6 domains. All proteins are expressed and purified from *E. coli*. “*” denotes bands corresponding to GST or GST-fused Snu71 domains. “+” denotes bands corresponding to Prp40 FF4-6 or FF4. Red labels indicate the GST-Snu71 domain (382-537) that stably interacts with Prp40 FF4 and FF4-6. **(C)** GST pull-down assay probing the interaction between Prp40 domains and BBP. Various GST-fused Prp40 domains as in (A) are used to pull down BBP. Pull-down samples are visualized on SDS PAGE with Coomassie stain. “*” denotes bands corresponding to various GST-fusion Prp40 domains (GST-FF2 and FF3 have significant degradation). “+” denotes bands corresponding to BBP. Red labels indicate Prp40 domains that pulled down BBP. **(D)** A summary of interactions among BBP, Prp40, Snu71, and Luc7 (indicated by solid line connecting regions of two proteins) as observed in panels A-C. **(E)** Key interaction regions mapped on the structures of Snu71, Prp40 (both are predicted by AlphaFold2 (Jumper et al. 2021)) and Luc7 (determined by cryoEM, (Li et al. 2019)). **(F)** A model of how Luc7, Snu71, and Prp40 interact with each other in the framework of U1 snRNP in the pre-A complex (Zhang et al. 2021).

### Luc7 depletion leads to mitochondrial dysfunction and galactose toxicity

In a serendipitous observation, we found that the growth medium of the Luc7D strain after Luc7 depletion consistently changed its color to brick red immediately after adding bleach for a safe disposal of the culture (**Fig. 5A**, top panel). Even the cell pellet turned to rusty red after prolonged induction (e.g., an overnight incubation with β-estradiol and auxin) (**Fig. 5A**, bottom panel). This indicated an intracellular as well as extracellular release of ferrous iron and its oxidation to red colored ferric iron by the bleach. We confirmed the release of iron using the iron-specific ferrozine assay (**Fig. 5B**). Since mitochondria are central for iron homeostasis in all eukaryotes, and many genes that code for mitochondrial proteins have introns, we examined splicing efficiency of these genes in the induced set. Nearly all intron-containing genes encoding proteins critical for mitochondrial biogenesis and function showed increased IR (**Fig. 5C, D**). Transcript abundance of these and that of the non-intronic genes with mitochondria-related function, as well as almost all genes that regulate iron homeostasis, was also affected to varying levels (**Supplementary Table 1**).

**Figure 5.**
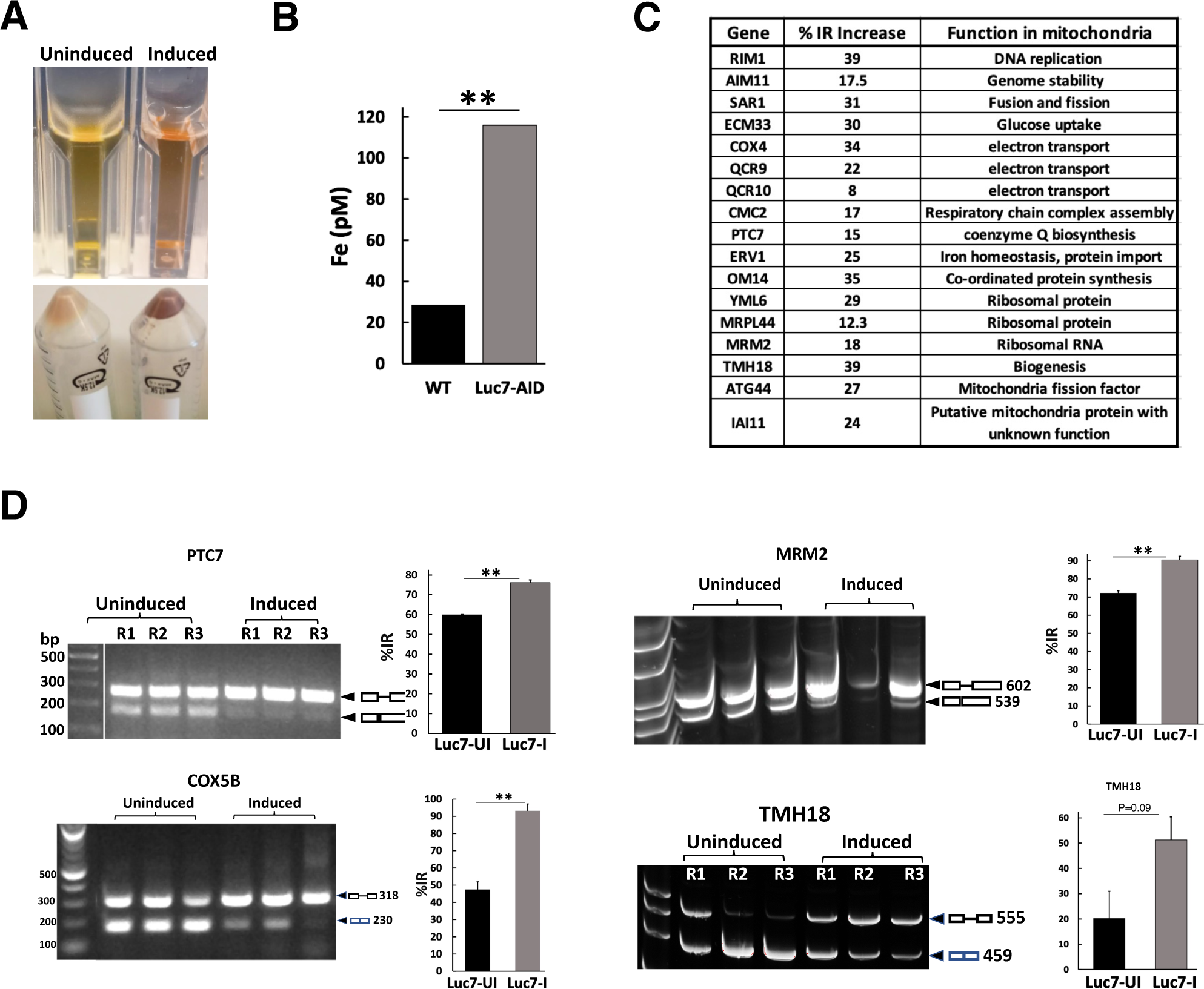
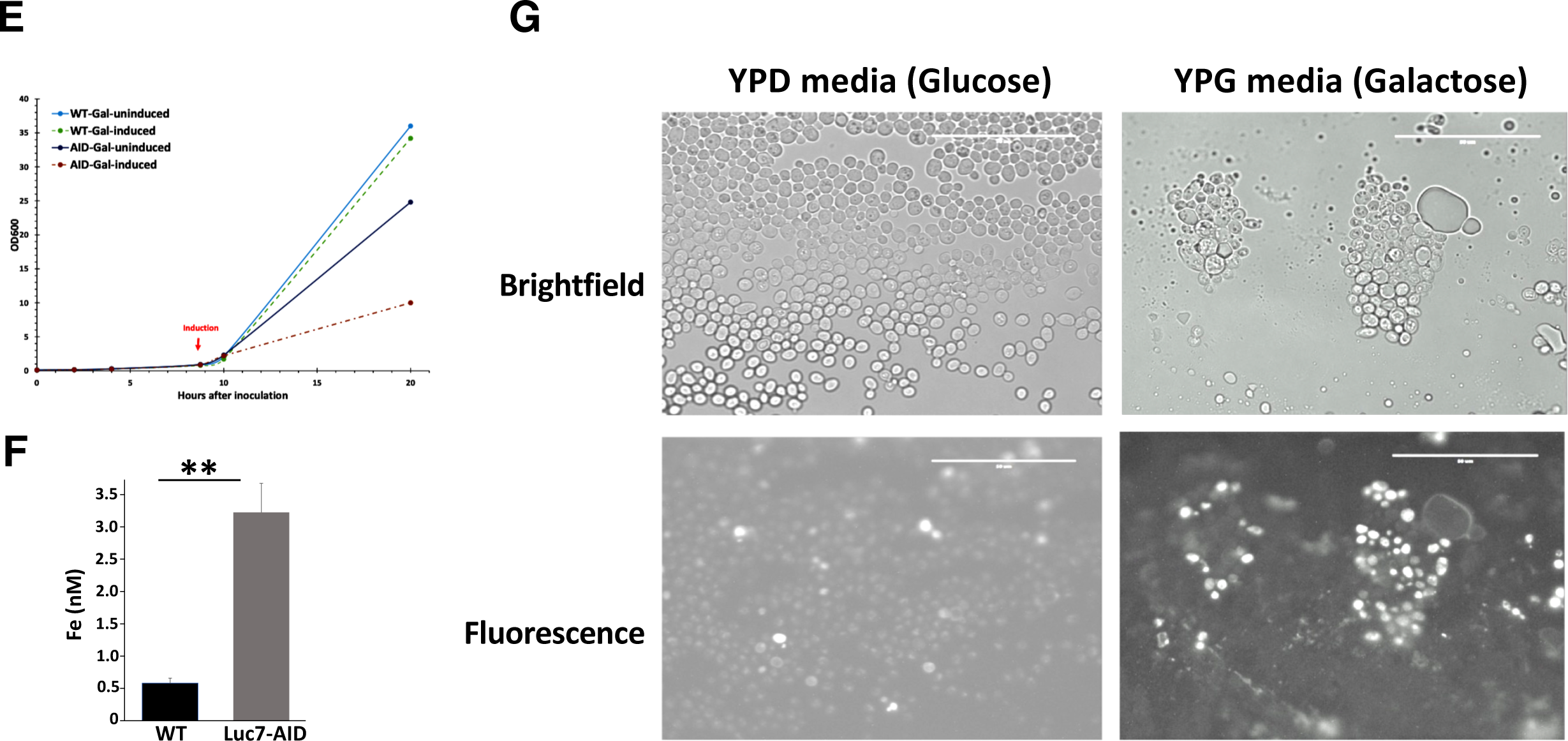
Luc7 depletion leads to mitochondrial dysfunction and galactose toxicity. **(A)** Luc7 depletion led to Fe^2+^ release. Both culture supernatants and cell pellets after induction turned rusty red, immediately after the addition of a few drops of 10% commercial bleach. (**B)** Free iron measurement using ferrozine assay in the WT and h-yU1 strain after overnight induction. (**C**) A list of pre-mRNAs with increased intron retention in RNAseq analyses of the Luc7D strain that affect mitochondrial integrity and function. IR: intron retention. (**D**) RT-PCR confirmation of RNA-seq results for three genes selected from the list in C and COX5B (which was not in the list of genes affected in the RNAseq data due to its extremely short exon (1 nt) which cannot be handled by typical sequencing analyses software). **(E)** Growth curves of auxin and β-estradiol (induced) or ethanol added (uninduced) Luc7D and WT yeast strains in galactose-medium (YPG). **(F)** Significantly more Fe^2+^ is released into the galactose medium of induced cultures (compared to panel B) as measured by ferrozine assay. **(G)** Dihydro Rhodamine detection of mitochondrial reactive oxygen species (which manifests as bright fluorescent spots within mitochondria in the fluorescent image) (Laun et al. 2001; Gomez et al. 2014) in the Luc7D strain grown in YPD- or YPG upon Luc7 depletion.

Although galactose is equally fermentable as glucose by budding yeast, early steps of galactose utilization require mitochondrial respiration. Galactose elevates heme biosynthesis in yeast, which in turn induces genes required for TCA cycle and OXPHOS (Zhang et al. 2017; De-Souza et al. 2020). Since certain mutations leading to mitochondrial dysfunction in mammalian cells are known to cause galactose toxicity (Iannetti et al. 2018), we compared the ability of Luc7D strain to utilize galactose under Luc7-sufficient versus Luc7-depleted conditions. As shown in **Fig. 5E**, the strain stopped growing in galactose soon after 6 h induction needed for Luc7 depletion and consequent impairment of mitochondrial function. The strain also released more Fe^2+^ when induced in the galactose medium than in the glucose medium (**Fig. 5F**, compare with **Fig. 5B**), suggesting that growth in galactose exacerbated the damage to mitochondria upon Luc7 depletion. This inference was further tested by staining Luc7D cells induced in glucose (YPD) or galactose (YPG) to detect mitochondria-localized reactive oxygen species (mtROS). As shown in **Fig. 5G**, a severe induction of mtROS was observed in YPG-grown cells when induced for Luc7 depletion.

## Discussion

U1C plays an important role in stabilizing the U1 snRNA and 5’ ss interaction. In structures of both the human and yeast U1 snRNP in complex with the 5’ ss RNA, the ZnF domain located in the N-terminal region of U1C contacts the backbone of the U1 snRNA and 5’ ss RNA duplex (Kondo et al. 2015). The ZnF domain of U1C is highly conserved between the two species, while their C-terminal domains are highly divergent (**Fig. 1A**). The C-terminal domain of yeast U1C is much longer and forms extensive interactions with Prp42, the latter facilitating the interaction of the other auxiliary proteins to the yeast U1 snRNP core (Li et al. 2017). Deletion of the CTD of U1C is lethal to yeast (Li et al. 2017). It may not be surprising that the full length human U1C cannot substitute yeast U1C (**Fig. 1B**), potentially because the much shorter and divergent human U1C CTD fails to interact with Prp42.

On the other hand, the ZnF domain of human U1C can functionally substitute that of yeast U1C, with no effect on yeast growth at 30 or 37 °C, and a slight growth retardation at lower temperatures (17 and 23 °C). The cryoEM structure of the U1 snRNP ^h-yU1C^ shows that the structure is essentially identical to the WT yeast U1 snRNP with the exception of different side chains in the U1C ZnF domain. Although the residues contacting the U1-5’ ss RNA duplex have minimal changes (T17 to S and R28 to K) from yeast to human, residues distant from the RNA-binding interface can potentially participate in long range allosteric signal transmission and affect the RNA-binding affinity, similar to what was observed in the case of U1A (Han et al. 2019). For example, a hydrophobic core composed of F4, L13, and the CH2 groups of R21 exists adjacent to the metal binding pocket in U1C and other canonical ZnF domains and stabilize the ZnF domain (Muto et al. 2004). Mutations at L13 reduces the interaction between U1 snRNA and the 5’ ss, bypassing the need for RNA helicase Prp28 that disrupts the U1 snRNA and 5’ ss interaction (Chen et al. 2001). In yeast U1C, F4 is replaced with a somewhat less hydrophobic Tyr residue, which can potentially have a similar effect as the L13 mutant, leading to reduced interaction between yeast U1 snRNA and 5’ ss than that of human. The Kd between U1 snRNP ^h-yU1C^ and 5’ ss is slightly lower than that between U1 snRNP ^WT^ and 5’ ss. It is possible that U1 snRNP ^h-yU1C^ binds 5’ ss tighter, leading to a reduced splicing efficiency. During the splicing cycle, U1 snRNP needs to bind and recognize the 5’ ss, but it also needs to be dissociated from 5’ ss (facilitated by Prp28) so it can be recognized by U6 snRNP (Staley and Guthrie 1999). It is plausible that the binding affinity between U1 and 5’ ss needs to be optimal and either too weak or too strong can lead to splicing defects. Given our hypothesis, one might expect that the genes with 5’ ss that form better complementation with the 5’ end of U1 snRNA will more likely have intron retention, although we have not observed significant correlation between increased intron retention and the distance to the yeast consensus 5’ ss or how well it complements the 5’ end of U1 snRNA (**Fig. 1L, M**). This is potentially because the ultimate splicing efficiency is determined by many other factors beyond just the 5’ ss strength. Genes with shorter intron or upstream exon seem to be more prone to intron retention in the h-yU1C strain (**Fig. 1N, O**), although the mechanism is yet to be understood.

In yeast U1 snRNP, a second protein Luc7 also uses its ZnF2 domain to help stabilizing the interaction between U1 snRNA and the 5’ ss (Plaschka et al. 2018; Li et al. 2019). The human homologs (Luc7L, L2, L3) of Luc7 are alternative splicing factors that only transiently associate with U1 snRNP and these proteins contain extra RS rich C-terminal domains that do not exist in yeast Luc7 (**Fig. 2A**). The yeast Luc7 could not be replaced with the full length human Luc7Ls (**Fig. 2B**). It is possible that the human Luc7Ls are different enough from yLuc7 such that they can no longer be incorporated into the yeast U1 snRNP, or the ZnF2 of hLuc7Ls is not sufficient for stabilizing the U1 – 5’ ss RNA duplex. Alternatively, the C-terminal RS domain of human Luc7Ls may have a dominant negative effect in yeast.

Previous cryoEM structures of the yeast E complex indicated that residues D212, R216, and H220 of Luc7 are in close proximity of the U1 snRNA and 5’ ss RNA duplex, however the density for Luc7 is poor and the side chains could not be modeled with high confidence. We generated the Luc7T triple mutant (D212A/R216A/H220C) and showed that U1 snRNP ^Luc7T^ has the same binding affinity to the 5’ ss and minimal splicing changes compared to the WT. These observations suggest that these three residues are unlikely to play a major role in the stabilization of the U1 snRNA and 5’ ss interaction.

Using an AID system, we were able to deplete Luc7 in the Luc7D strain (which was combined with the humanized ZnF domain of yU1C). Although Luc7 is an essential gene, the Luc7D strain continued to grow even after prolonged auxin induction (**Fig. 3B**). This may be due to a residual amount of U1 snRNP containing Luc7 remained in cells, even though the majority of U1 snRNP is depleted of Luc7 after 6 h induction (**Fig. 3D**). U1 snRNP without Luc7 can be assembled, although it also completely lost Snu71 and Prp40 and is less stable than the WT U1 snRNP. This is consistent with previous observations that a Luc7 mutant leads to the reduced association of Snu71 and Prp40 with U1 snRNP (Fortes et al. 1999). By consolidating previous cryo-EM structures, crosslinking and mass spectrometry analyses, and pull-down analyses presented here, we present a model of how these three proteins interact with each other and the rest of the U1 snRNP (**Fig. 4E, F**). In essence, Luc7 binds to the Sm ring through its N-terminal helix and ZnF1, to the U1 – 5’ ss RNA duplex through its ZnF2, and to Snu71 through its CC domain. Snu71 binds Prp42 and Snu56 in U1 snRNP through its N-terminal domain (residues 1-50), and to Luc7 through the long helix formed between residues 260-312 (Zhang et al. 2021). Prp40 binds to the C-terminal domain of U1-70K through its FF6 domain (Li et al. 2019), to a region within residues 382 to 537 of Snu71 through its FF4 domain (this paper) and potentially FF2 and 3 domains (Zhang et al. 2021), and to BBP through its N-terminal WW domain. Although both Snu71 (through its N-terminal domain) and Prp40 (through FF6) contact U1 snRNP, the interaction between Luc7 and U1 snRNP seems to be critical for maintaining the trimer within the U1 snRNP, as evident from the loss of Prp40 and Snu71 when Luc7 is depleted (**Fig. 3D, E**). Given that Prp40 is critical for bridging the 5’ ss and BPS (Li et al. 2019; Zhang et al. 2021), it is not surprising that Luc7 depletion (and the subsequent loss of Prp40 and Snu71 from U1 snRNP) led to extensive splicing defects in essentially all intron-containing yeast genes with sufficient reads in RNAseq analysis.

The vast majority of yeast intron-containing genes encode ribosomal proteins, so studies of the functional consequence of widespread splicing defects have usually focused on ribosomal proteins. We showed that another important consequence of substantial splicing defects in yeast is mitochondria dysfunction. Splicing of essentially all intron-containing gene related to mitochondria function are negatively impacted (**Fig. 5C**). The resulting mitochondria dysfunction likely led to an impaired iron homeostasis and galactose toxicity. Interestingly, human Luc7L2 KD led to significant downregulation of glycolytic genes (Daniels et al. 2021), and Luc7L2 was found to be a major player in maintaining the balance between oxidative phosphorylation (OXPHOS) in mitochondria and glycolysis in cytosol, the two major pathways for ATP production (Jourdain et al. 2021). Luc7L2 suppresses OXPHOS through alternative splicing regulation of specific genes (the glycolytic enzyme PFKM and cystine/glutamate antiporter SLC7A11) as well as secondary repression of mitochondrial respiratory complex protein expression. Although the mechanism of regulation of energy metabolism by yeast Luc7 (through global suppression of splicing which in turn affects essentially all intron-containing genes encoding mitochondrial proteins) and human Luc7L2 (through specific alternative splicing regulation of PFKM whose yeast homology PFK2 is intronless and SLC7A11 which has no yeast homolog) is apparently different, it highlights the important and conserved role of splicing in the regulation of eukaryotic energy production.

## Materials and Methods

### Yeast strains and growth

All yeast strains used were grown on YPD or appropriate dropout media. 50 µg/mL of ampicillin was routinely incorporated into large scale liquid cultures to avoid bacterial contamination. Plasmid shuffling was done by selecting initial transformants (from dropout plates) on 5- fluorourotic acid (5-FOA). Colonies resistant to 5-FOA were retested on appropriate dropout plates to confirm successful shuffling.

### Humanizing yeast U1C

The first 36 residues of yeast U1C protein encompassing the zinc-finger domain were replaced with the corresponding region of the human U1C protein using the CRISPR-Cas9 system in a yeast Luc7 shuffle strain (Agarwal et al. 2016), essentially as described in (Laughery and Wyrick 2019). The following three guide RNAs covering the first 108 bp of the coding sequence were cloned into pML107 (Laughery and Wyrick 2019) and used to swap the first 36 amino acid segment with that of the human U1C: 5’-ATGACACGTTGAGCGTT-3’, 5’-GTTCGTAAATCGCACTTGG-3’, 5’- CGTATAACAGCTGACTATT-3’.

The yeast-codon optimized human sequence (lower case indicates changed bases) with flanking sequences from the yeast Luc7 gene (italics), as shown below, was cloned into pUC19 by Gibson cloning (Hi-Fi cloning system, NEB) to replace the yeast sequence:

5’- *tagtgtaggcgatgaaggtgctcaagtaacggagaggaaagagataggca*ATGCCaAAaTTTTATTGTGAtTAtT GtGATACtTAttTgACtCATGAtTCTCCATCTGTtAGAAAaACtCAtTGttcTGGAAGaAAACAt AAAGAaAATGTtAAAGA*ttattatagAaacaaagcaagagacattattaataaacataat*-3’.

### Generating an inducible Luc7 depletion strain

We introduced an inducible degradation system for yLuc7 in the humanized U1C (h-yU1C) strin (described above). We engineered an auxin-inducible degron (AID) + 3x-HA tag at the C-terminus of yLuc7 and an auxin receptor (OsTIR1) driven by a β-estradiol inducible promoter (Mendoza-Ochoa et al. 2018) cloned into a single plasmid (pRS415) and transformed it into the h-yU1C strain.

### Phenotypic Analyses of Yeast

OD600 was used to monitor the growth in YPD (glucose as the sugar) or YPG (galactose as the sugar). Spot assay was done on YPD plates with a starting OD600 of 0.5 and 4x serial dilution. Fe^2+^ was measured using ferrozine, an iron-specific reagent (Riemer et al. 2004). Yeast cells were stained for viability using Evans Blue and mitochondrial ROS using dihydrorhodamine (Gomez et al. 2014). All microscopy work was done using EVOSfl all-in-one Olympus microscope (Advanced Microscopy Group, Bothell, WA) as described (Denney et al. 2021).

### Total protein isolation and immunoblot analysis

Total cellular protein from mid-log phase grown yeast was extracted using the TCA precipitation protocol (Yaffe and Schatz 1984) as modified by (Denney et al. 2021). The protocol preserves the intactness of proteins by suppressing proteolysis and any post-translational modifications. Equal amount of protein quantified as A280 was resolved by PAGE (10% acrylamide-SDS), blotted onto a PVDF membrane and probed with α-HA tag (Roche, Catalog# 11867431001; for the detection of AID and HA-tagged Luc7) and α-CBP tag (GenScript, catalog # A00635; for the detection of U1A) antibodies. The immune-reactive proteins were detected using HRP-labeled secondary antibodies and chemiluminescence.

### Purification of U1 snRNP ^WT, h-yU1C, Luc7T, Luc7D^

Purification of U1 snRNP ^WT, h-yU1C, Luc7T^ were carried out essentially as described (Li et al. 2017). U1 snRNP ^Luc7D^ was purified from yeast cells by growing them in 2xYPD to OD600 of 8-10 and inducing them for 6 h by adding 10 µM of estradiol and 750 µM of IAA.

To analyze U1 snRNP level in cells, 5 ml of U1 snRNP ^WT, h-yU1C, Luc7T^ were grown at 30 °C or 17 °C and purified by one step IgG purification. Whole cell and IgG elution were analyzed by Western blot with an anti-CBP antibody and solution hybridization with an IRDye-700 labeled U1-specific primer.

Purified U1 snRNP ^h-yU1C^ were assembled with ACT1 pre-mRNA and BBP-Mud2 essentially as described (Li et al. 2019). In brief, ACT1 pre-mRNA bound to MBP-MS2 fusion protein was mixed with IgG Sepharose-6 purified h-yU1C, BBP-Mud2 dimer and calmodulin resin (Agilent). After incubation at 4°C for 3 h, the resin was washed 8 times, and then eluted six times with 100 µl eluting buffer. The elutions were pooled for cryo-EM sample preparation.

### Fluorescence Polarization Assay

Serial dilutions (1:2) of each U1snRNPs were prepared in a buffer containing 20 mM HEPES 7.9, 150 mM KCl, 2 mM EDTA, 1 mM MgCl2, 0.01% Triton X-100 and 0.1% PEG3350. Cy5-labeled RNA oligo (GAUUCUG<u>GUAUGU</u>UC, from Integrated DNA Technologies) containing the canonical 5’ ss (underlined) was added into each sample at a final concentration of 3 nM. The mixture was transferred into a 384-well, black, flat-bottom microplates (Greiner Bio-One) in duplicates and were incubated at room temperature for 15min. The plate read using an Envision plate reader (PerkinElmer) at excitation/emission = 620/688 nm) at 25°C. The experimental data were analyzed using Prism 10 software (Graphpad Software, San Diego, CA, USA).

### Cryo-EM sample preparation and imaging

Three microliters of the freshly purified complex were applied to a plasma-cleaned C-flat holy carbon grid (1.2/1.3, 400 mesh, Electron Microscopy Sciences) with 20 s glow discharge using air and flash-frozen in liquid ethane with a Vitrobot Mark IV (Thermo Fisher Scientific). The grids were obtained with the chamber at 100% humidity, 2.5 s blotting time, –6 blotting force and 15 s wait time and flash-frozen into liquid ethane with a Vitrobot Mark IV (Thermo Fisher Scientific). The data was collected on a Titan Krios at the Pacific Northwest Cryo-EM Center (PNCC) operated at 300 keV and equipped with a K3 direct detector (Gatan). Movies were acquired at a pixel size of 0.83A/pix (0.415A/pix super-res), a defocus range of −0.8 to −2.5 μm, and 50 frames with a total dose of ∼50 e-/Å^2^.

### Image processing, model building, and refinement

Patch motion correction and patch CTF estimation were done in cryoSPARC. A total of ∼1.5 million particles were automatically picked using “Blob Picker” and extracted with a box size of 300 × 300 pixels. After three rounds of reference-free 2D classification, ∼71,607 particles were selected for ab initio reconstruction, and heterogeneous refinement. 138,997 good particles are then unbinned with a box size of 640 × 640 pixels. After homogenesis refinement and nonuniform refinement, the map reached a resolution of ∼3.49 Å.

The model was built in Coot. To aid subunit assignment and model building, we took advantage of the reported E structure (PDB code: 6N7R) which was fitted into the h-yU1C density map. The first 36 aa of U1C and the 5’ss region were adjusted manually to match the density. The RNA components were subsequently adjusted using RCrane. The rest of the subunits fit well and are slightly adjusted for the side chains. The model was refined using PHENIX in real space with secondary structure and geometry restraints.

### RNA-seq Analysis

Total RNA was isolated from the yeast strain with humanized U1 grown in YPD in triplicates of 4 h-uninduced and 6 h-induced treatments. Hot phenol (Green and Sambrook, 2021) extracted RNA was DNase I-treated and purified further using Monarch RNA Cleanup spin-columns (NEB). Library preparation and short-read sequencing was done by Novogene. Reads were mapped to the yeast reference genome (version R64.3.1), using STAR program (Dobin et al. 2013). Differential expression analysis was carried out using DE-seq2 (Love et al. 2014) and splicing analysis using JunctionCounts (github ajw2329/junctionCounts) programs.

Differentially included introns were identified by comparing PSI values across conditions using a Benjamini-Hochberg corrected t-test. Affected introns were defined as those with a p value of less than 0.05 and an absolute difference in mean PSI value across conditions of at least 0.05. Distances of affected 5’ splice sites to the consensus yeast 5’ splice site (GTATGT) were calculated using Hamming distances.

### RT-PCR analysis

Total RNA, purified as described above, was used in sequential reverse transcription and PCR amplification steps using the Protoscript II first strand cDNA synthesis kit and Q5 Hi-fidelity DNA polymerase (NEB) according to manufacturer’s protocols. 1-2 µg of total RNA was used for reverse transcription and 1-2 µL of the reaction product was amplified by PCR. Primer sets that can amplify both the unspliced and spliced forms of selected mRNA were used in each case. Appropriate number of amplification cycles was chosen for each amplicon, based on the copy number/cell data (available from SGD) and as needed for optimal quantitation of the products from agarose or acrylamide gel images. Signals were quantified using Image J program (https://imagej.net/software/imagej/).

### Pull-down assay from yeast after co-transformation

The coding regions of yeast Prp40, Snu71, and Luc7 full length or truncations were amplified by PCR using genomic *S. cerevisiae* DNA as templates, and ligated into pRS414, pRS416 and pRS317 vectors. The final plasmids constructed are: pRS414/GPD-protA-Prp40, pRS317/GPD-Prp40, pRS416/GDP-Snu71-CBP, pRS414/GPD-His6-Luc7 and pRS414/GPD-His6-Luc7ΔCC (Luc7 coiled coil domain (residues 123-190) deletion). Yeast BCY123 cells were transformed with different combination of the plasmids (as indicated in **Fig. 4A, B**) and selected on appropriate selective media. Clones from the transformation were cultured in 50 mL of liquid selective media to OD600=3-4. Cells were harvested and lysed in lysis buffer (50 mM Tris-HCl (pH 8.0), 150 mM NaCl, 0.05% NP40, 1 mM DTT, 2.5mM CaCl2, 1.5mM MgCl2) using the bead-beating method. The lysates were applied to either calmodulin resin (if CBP tagged Snu71 is expressed) or IgG resin (if protA tagged Prp40 is expressed). The resins were washed with the lysis buffer. The proteins were eluted off calmodulin resin using the eluting buffer (20 mM Tris-HCl (pH 8.0), 150 mM NaCl, 0.01% NP40, 2.5mM EGTA, 1 mM MgCl2, 0.5mM DTT) or cleaved off IgG resin using TEV protease in the same buffer. The proteins were separated on SDS-PAGE and transferred to a nitrocellulose membrane. Western blot was performed using an anti-His tag antibody.

### GST pull-down assays using purified proteins

Snu71 fused to a C-terminal CBP tag was cloned into the plasmid pRS416/GPD-Mud2-CBP. Snu71 was purified from yeast BCY123 cells harboring the above plasmid using calmodulin resin. BBP was purified from BCY123 cells harboring pRS414/GPD-protA-BBP using IgG Sepharose-6 Fast Flow resin. Various Prp40 domains fused with an N-terminal Flag tag were cloned into pGEX-6p-1 vector (GE Healthcare) and purified from *E.coli* as GST fusion proteins. The Prp40 domains constructed are: 1-75 (WW); 134–189 (FF1); 198–259 (FF2); 260–335 (FF3); 351-415 (FF4); 425–488 (FF5); 488-552 (FF6); 351–552 (FF4-6). To detect interactions between Prp40 domains with Snu71 or BBP, GST fusion proteins or GST alone were bound on glutathione-Sepharose resin (GE Healthcare) and incubated with 5 μg of purified Snu71 or BBP protein in pull-down buffer (20 mM Tris pH 8.0; 120 mM NaCl, 1mM Mg2Cl, 1 mM DTT; 0.05 % Triton-X-100) for 2 h at 4°C. Resins with bound proteins were washed five times with pulldown buffer. Resins were boiled in SDS loading buffer and proteins were analyzed on SDS-PAGE followed by Coomassie stain.

To detect interactions between Prp40 FF domains with different regions of Snu71, GST fusion of FF4, FF5, FF6 and FF4-6 were purified using glutathione-Sepharose resin and released by PreScission protease. The proteins were further purified by gel filtration using a Superose 6 Increase 10/300 GL column (GE Healthcare). Snu71 fragments with residues 1-96, 97-220, 221-380, 381-537, 537-620 fused with an N-terminal HA tag were cloned into pGEX-6p-1 vector. Snu71 fragments 1-96 and 221-381 failed to be expressed and purified. GST fusion of the other Snu71 fragments or GST alone were coupled to glutathione-Sepharose resin and incubated with 2ug of each individual purified FF domains in pull-down buffer (20 mM Tris pH 8.0; 120 mM NaCl, 1mM Mg2Cl, 1 mM DTT; 0.05 % Triton-X-100;) for 2 h at 4°C under rotation. Resins with bound proteins were washed five times with pulldown buffer. Resins were boiled in SDS loading buffer and proteins were analyzed on SDS-PAGE followed by Coomassie stain and western blot. Western blot was performed using M2 antibody (Sigma-Aldrich).

### Mass Spectrometry

Samples were subjected to proteolytic digestion using a filter-aided sample preparation (FASP) protocol (Wisniewski et al. 2009) with 10 kDa molecular weight cutoff filters (Sartorius Vivacon 500 #VN01H02). Samples were reduced with 5 mM tris(2-carboxyethyl)phosphine), alkylated with 50 mM 2-chloroacetamide, and digested overnight with trypsin (enzyme:substrate ratio 1:50) at 37°C. Peptides were recovered from the filter using successive washes with 0.2% formic acid (FA). Aliquots containing 10 μg of digested peptides were cleaned using Pierce^TM^ C18 Spin Tips (Thermo Scientific) according to the manufacturer’s protocol, dried in a vacuum centrifuge, and resuspended in 0.1% FA in mass spectrometry-grade water.

Liquid chromatography-tandem mass spectrometry (LC-MS/MS) was performed using an Easy nLC 1200 instrument coupled to an Orbitrap Fusion Lumos Tribrid mass spectrometer (all from ThermoFisher Scientific). Tryptic peptides were loaded on a C18 column (100 μM inner diameter x 20 cm) packed in-house with 2.7 μm Cortecs C18 resin and separated at a flow rate of 0.4 μl/min with solution A (0.1% FA) and solution B (0.1% FA in ACN) and under the following conditions: isocratic at 6% B for 3 minutes, followed by 6%-42% B for 102 minutes, 42%-60% B for 5 minutes, 60%-95% B for 1 min and isocratic at 95% B for 9 minutes. MS/ MS was performed using data-dependent acquisition (DDA) mode with a cycle time of 3 s between master scans and dynamic exclusion was set to 45 s. Fragmentation spectra were interpreted against the UniProt *Saccharomyces cerevisiae* proteome database (Proteome ID # UP000002311) using the MSFragger-based FragPipe computational platform (Kong et al. 2017). Contaminants and reverse decoys were added to the database automatically. The precursor-ion mass tolerance and fragment-ion mass tolerance were set to 10 ppm and .2 Da, respectively. Fixed modifications were set as carbamidomethyl (C), variable modifications were set as oxidation (M) and two missed tryptic cleavages were allowed. The protein-level false discovery rate (FDR) was ≤ 1%.

## Acknowledgements

This work was supported by NIH grants R35GM145289 and R01GM126157 (R.Z.); R35GM133385 (J.M.T.); R35GM147498 (J.W.); and R01GM071940 (Z.H.Z.). J.G. was supported by the NIH under Ruth L. Kirschstein National Research Service Award T32CA17468. The contents are solely the responsibility of the authors and do not necessarily represent the official views of the NIH. We thank Dr. Manny Ares (University of California, Santa Cruz) for advice on yeast pre-mRNA RNA splicing analysis using short-read sequencing, Drs. Beate Schwer and Stewart Shuman (Weill Cornell Medical College, Cornell University) for the gift of U1C and Luc7 shuffle strains, and Dr. Michael McMurray (Department of Cell & Developmental Biology, University of Colorado Anschutz Medical Campus) for advice on yeast genome-editing as well as the use of EVOSfl microscope for yeast imaging and Kate Matlin for help with RNAseq data mining. A portion of this research was supported by NIH grant U24GM129547 and performed at the PNCC at OHSU and accessed through EMSL (grid.436923.9), a DOE Office of Science User Facility sponsored by the Office of Biological and Environmental Research. We acknowledge the staff (in particular Rose Marie Haynes) at the Pacific Northwest Cryo-EM Center for help with data collection. We also acknowledge support from the CU Anschutz School of Medicine Cryo-EM core, the proteomics core, and the HTS Drug Discovery and Chemical Biology core facilities (partially supported by the School of Medicine and the University of Colorado Cancer Center Support Grant P30CA046934). Molecular graphics and analyses were performed with the UCSF Chimera and ChimeraX, developed by the Resource for Biocomputing, Visualization, and Informatics at the University of California, San Francisco, with support from NIGMS P41-GM103311 (Chimera, ChimeraX) and NIH R01-GM129325 (ChimeraX).

---

**Supplementary Figure 1.**
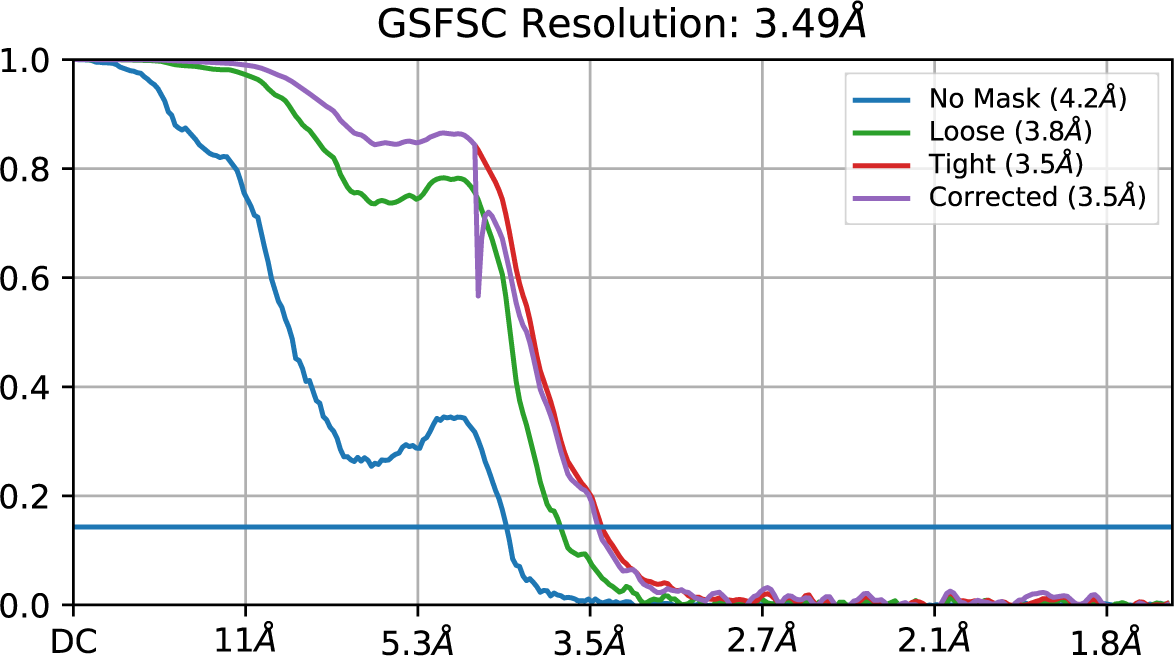
The gold-standard FSC curve of the final reconstruction indicates a numerical resolution of 3.5 Å.

## Notes

### Competing Interest Statement

The authors have declared no competing interest.

